# The sensitising effect of IgG in fibromyalgia syndrome is mediated by Mrgprb2 in mast cells

**DOI:** 10.1101/2025.05.15.652596

**Authors:** Karla R. Sanchez, Jamie Burgess, Qin Zheng, Uazman Alam, Harvey Neiland, Richard Berwick, David Andersson, Samantha Korver, Anne Marshall, Andreas Goebel, Xinzhong Dong

## Abstract

Fibromyalgia syndrome (FMS) is characterized by elevated levels of immunoglobulin G (IgG), altered bowel habits, and increased pain sensitivity, suggesting immune dysregulation, but the exact mechanism remains unclear. Here, we found that FMS-IgG binds to mast cells in a MRGPRX2/b2-dependent manner, leading to mast cell recruitment and IL-6 secretion. Transferring serum-IgG from FMS patients to mice induced FMS-like symptoms and increased skin mast cells, indicating that FMS-IgG acts through mast cell activation. The ablation of mice Mrgprb2 mast cells or deleting Mrgprb2 receptors prevented IgG-induced heightened sensitivity to mechanical and cold stimuli. Stimulating human LAD2 cells with FMS IgG elicited MRGPRX2-dependent IL-6 production. Consistent with mice findings, mast cell density and tryptase levels increased in human FMS skin samples compared to healthy controls. Taken together our results suggests that FMS IgG mediates hypersensitivity via activation of mast cells bearing the MRGPRX2 receptor and that these cells are a potential therapeutic target.

## INTRODUCTION

Fibromyalgia syndrome (FMS) is a chronic condition characterised by widespread musculoskeletal pain, fatigue, sleep disruptions, and cognitive impairment^1^. Individuals with severe forms of this disorder often struggle to maintain activities of daily living, leading to significant personal distress and substantial economic burden including for families and the economy^2, 3^. Despite the high prevalence of FMS, therapeutic options remain limited. While many patients may experience modest improvements with light exercise, recommended medication treatment with low-dose tricyclic antidepressants and serotonin-norepinephrine reuptake inhibitors is effective in only a minority of patients and often drug tolerance develops^4^. This highlights the critical need for new, effective therapeutic strategies informed by a deeper understanding of the underlying pathophysiology of FMS. Recent evidence indicates that many patients with severe FMS harbour immunoglobulin G (IgG) autoantibodies that may be responsible for the core symptoms of the disorder^5^. Purified serum IgG from FMS patients (FMS-IgG) injected into rodents induces profound mechanical and thermal hypersensitivities, reduced locomotion, and decreased grip strength^6^. Additionally, sensory nerve fibres, including C fibres and Aδ fibres, show increased responsiveness to mechanical stimuli, and the small nerve fibre density in the skin is reduced^7^. Notably, IgG from healthy controls (HC; HC-IgG) and FMS serum from which IgG has been depleted do not produce these effects. These findings meet the Witebsky-Rose criteria for autoimmunity, suggesting that mechanisms of FMS may be underpinned by IgG autoimmunity^8^. Increased binding of patient-derived IgG to satellite glial cells surrounding sensory nerve bodies in the dorsal root ganglia (DRG) correlates with the pain intensity reported by FMS patients^8^. However, the mechanisms through which IgG causes FMS symptoms have remained unclear. Better understanding could pave the way for therapeutic strategies that go beyond reducing serum immunoglobulin levels.

A growing area of interest in FMS research is the involvement of mast cells in pain modulation^9, 10^. Mast cells, traditionally associated with allergic responses, are located near peripheral nerves and can influence pain sensitivity through the release of pro-inflammatory mediators such as histamine, cytokines (IL-6), and proteases (tryptase)^9^. Certain genetic variants involved in mast cell signalling and degranulation have been suggested associated with FMS^11, 12^. Prior research has demonstrated an increased density of skin mast cells in FMS, suggesting a potential link between mast cell activation and clinical symptoms^13, 14^.

In both humans and rodents, specific receptors expressed on mast cells, such as MRGPRX2 (humans) and Mrgprb2 (rodents), play a crucial role in mast cell activation^15, 16^. These receptors, which respond to a broad range of ligands, trigger mast cell degranulation, releasing substances including cytokines (IL-6) and chemokines (CCL2) to amplify pain signals^9,17^and promote further mast cell recruitment^18, 19^. IgG has not, been reported to directly bind to these receptors, however it can bind to Fc receptors (FcγRs) on mast cells. promoting degranulation with release of pro-inflammatory mediators such as histamine, cytokines (IL-6), and tryptase which can contribute to nociceptor sensitisation ^9, 20–22^. These processes cause hyperalgesia and allodynia, where painful stimuli are perceived as more painful, and normal stimuli are perceived as painful respectively^9, 23, 24^. Antidromic release of substance P from already-hyperactive sensory fibres activates mast cells to drive the release of mediators and further sensitise these fibres, perpetuating the pain phenotype^25^. We hypothesised that pro-algetic FMS IgG exerts its effect through activation of mast cells.

## RESULTS

### Administration of human FMS-IgG in mice induces pain-related behaviour

To explore the role of mast cells in the development of FMS-typical hypersensitivity to stimuli, we administered human FMS-IgG systemically to wild type (WT) mice and observed its effects on pain behaviour and mast cell activation (Fig. 1A). Mechanical hypersensitivity was assessed using Von Frey filament test, a widely accepted method for quantifying pain sensitivity in response to mechanical stimuli. Mice were tested by applying calibrated filaments with forces 0.07 g and 0.4g to the plantar surface of both hind paws. Mice treated with FMS-IgG exhibited a significant decrease in paw withdrawal thresholds by Day 4 with each of three FMS preparations, indicating heightened mechanical sensitivity compared to mice treated with HC-IgG (Fig.1 B-C). We utilised the Cold Plantar Assay to evaluate cold allodynia, measuring paw withdrawal latencies in response to a cold stimulus applied to the plantar surface (Fig.1D). Mice treated with FMS-IgG demonstrated a significant decrease in paw withdrawal latencies compared to HC-IgG treated mice (Fig. 1D).

**Figure 1.**
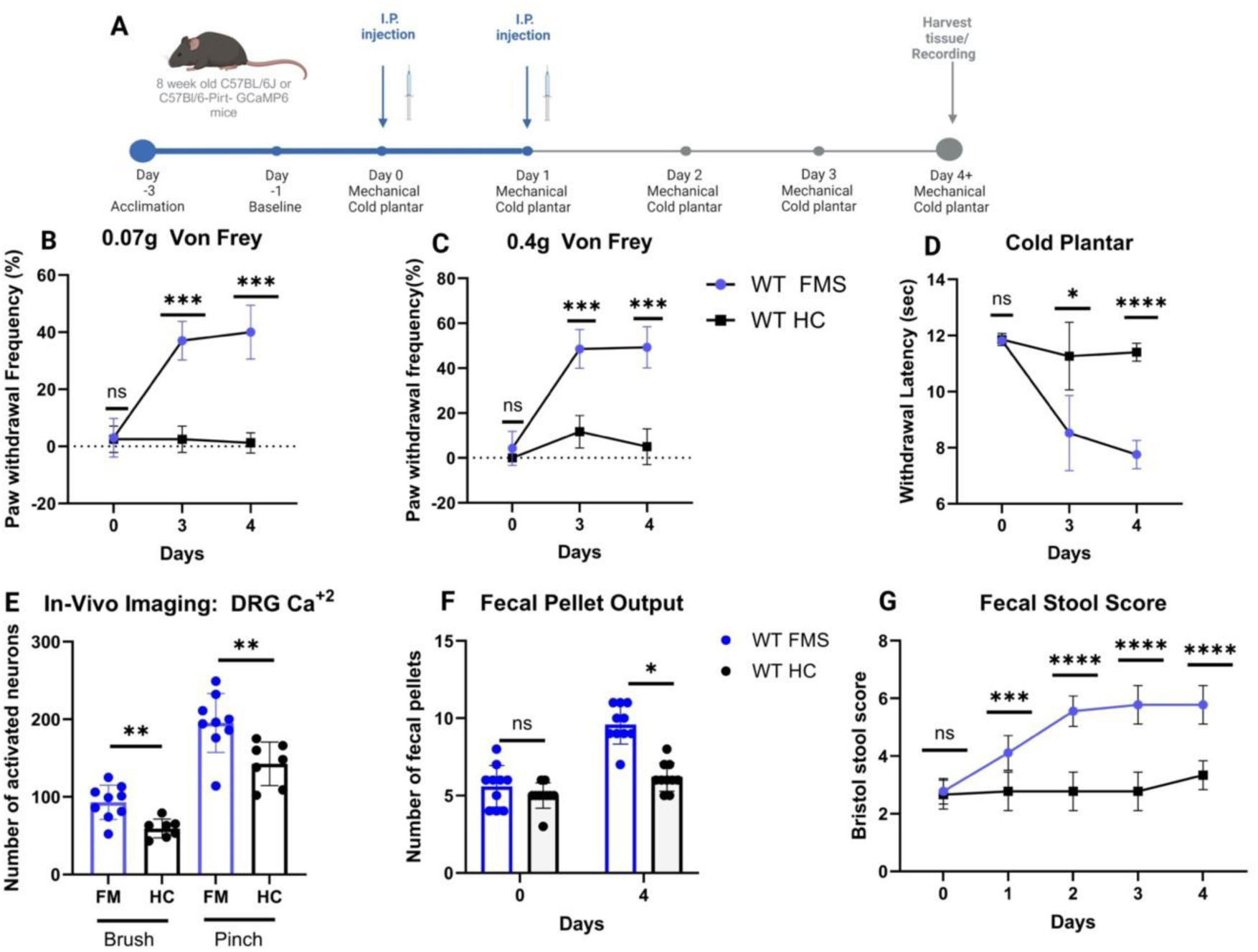
Administration of human fibromyalgia immunoglobulin (IgG) in mice induces pain behaviour and increased faecal output. Mice received intraperitoneal injections (IP) of 8mg IgG from either of 3 different fibromyalgia patients or healthy controls over 2 consecutive days to evaluate whether IgG produces hypersensitivity. (A) Schematic of the experimental design, for live behavioural recording or tissue collection. (B-C) Mechanical sensitivity was assessed using von Frey test at low (0.07g) and high (0.4g) forces, from both hind paws. (D) Cold allodynia was measured using the cold plantar assay, and responses from both hind paws collected. Data represent behavioural nociception assay in FMS and healthy control (HC) groups. (E) *In vivo* imaging of L4 DRG quantified the neurons responding to stimuli (individual dots represent mice). (F-G) Stool production and classification were quantified to evaluate gastrointestinal effects. Statistical significance is indicated as ns = not significant, *p < 0.05, **p < 0.01, ***p < 0.001, and ****p < 0.0001 for FMS vs. HC. Data are presented as mean ± SEM (n = 10/group for behaviour testing; n = 7-8/group for in vivo imaging) and were analysed using one- or two-way ANOVA followed by Bonferroni post hoc test. Nonparametric data were expressed as median and analysed using the Kruskal-Wallis test followed by Dunn’s test. Created with BioRender.com.

To assess the impact of FMS-IgG on neuronal activity, we performed in-vivo calcium (Ca²⁺) imaging on L4 dorsal root ganglion (DRG) neurons in live Pirt-Cre GCaMP6 mice. This genetically modified model allows for real-time visualization of neuronal activity. The technique provides insights into the excitability of DRG neurons in response to various stimuli, helping to assess changes in sensory neuron activity, particularly in pain-related studies. For this experiment, a noxious, high intensity pinch and a dynamic, low intensity brush stimuli were applied. Following FMS-IgG administration, DRG neurons exhibited significantly higher calcium influx in FMS mice compared to HC-IgG-treated controls in response to either stimulus, indicating enhanced neuronal excitability (Fig. 1E). These findings strongly indicate that passive treatment with IgG from FMS patients induces nociceptor activation compared to IgG from HC (HC-IgG).

An unexpected increase in faecal output and softer stool consistency was observed in mice treated with all three preparations of FMS-IgG, not reported previously; these changes are suggestive of irritable bowel syndrome (IBS)-like bowel irregularities commonly reported by people with FMS (Fig. 1F-G).

### Treatment with human FMS-IgG increases mast cell numbers in hind paws and back skin of wildtype mice

To investigate whether the mast cells skin density was increased, 14µm sections of the glabrous hind paw footpad were harvested from samples taken from all three FMS-IgG injected animal groups (n = 9 experiments/treatment) on day 4 and stained using immunofluorescence. There was a three-fold increase in Avidin positive mast cells (Fig. 2A) after treatment with FMS-IgG relative to HC-IgG (p < 0.001) (Fig. 2B). Significantly increased mast cell density at day 4 was also identified in 4µm wax preparations of non-glabrous dorsal mouse skin (n=1/treatment) assessed via brightfield microscopy (Fig. S4).

**Figure 2.**
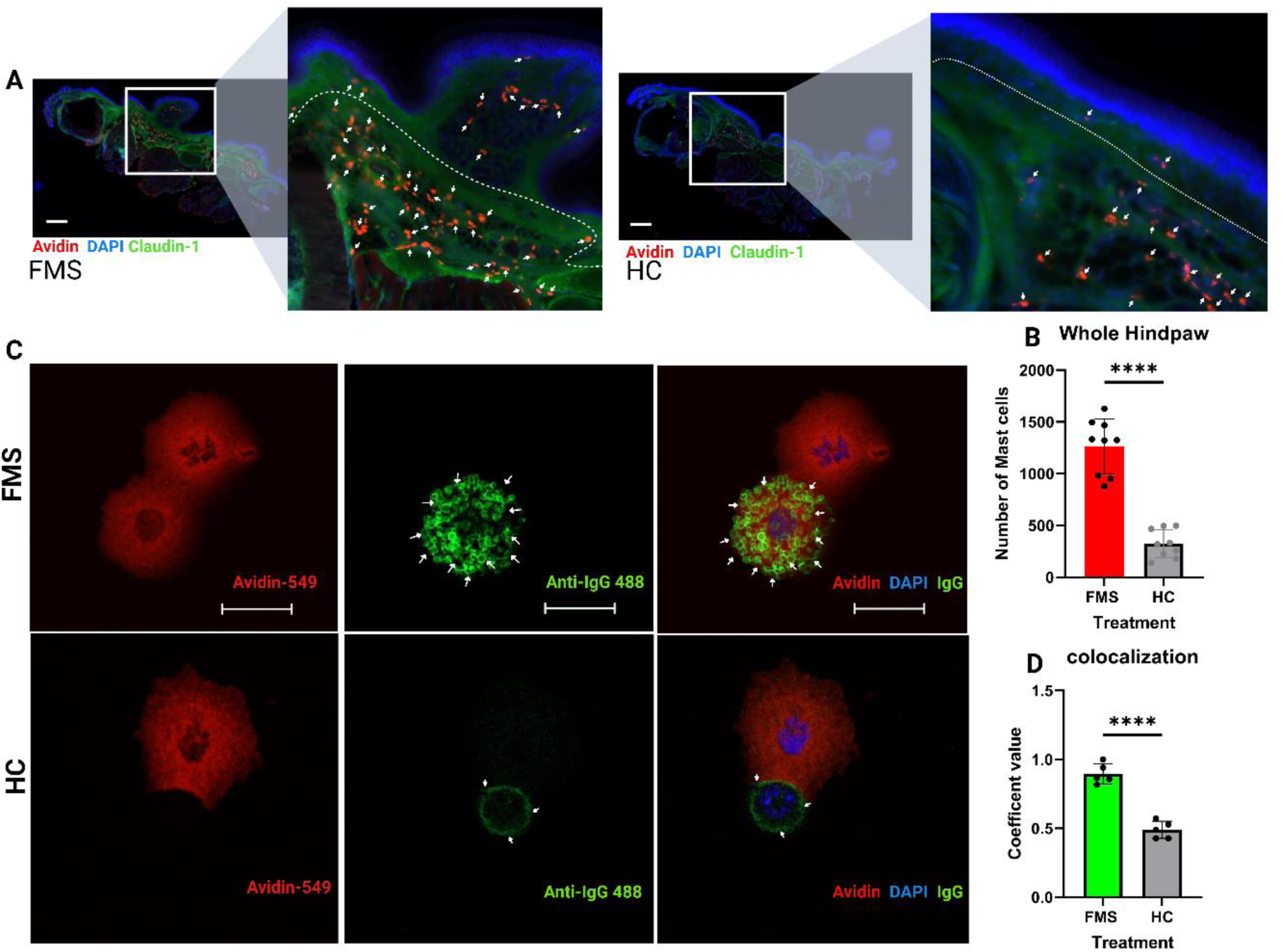
Treatment with human FMS-IgG significantly increases mast cell numbers in the hind paws of wildtype mice, and IgG binding in their peritoneal mast cells. (A-B) Double immunofluorescence was performed on 14 µm sections of the glabrous hind paw harvested on day 4. Mast cells were labelled in red (avidin), claudin-1 in green, and nuclei in blue (DAPI). White arrows indicate mast cell distribution between treatments. Images are zoomed in to match the same orientation and size; scale bar = 100 µm. Avidin-positive cells in the dermis were counted manually using confocal microscopy, and data represent at least 9 separate animals/treatment. (C-D) Peritoneal mast cells (PMCs) were collected on day 4 post-treatment and concentrated using Cytospin. Mast cells were stained in red (avidin), human IgG in green (anti-IgG 488), and nuclei in blue (DAPI). White arrows point to IgG-positive granules on mast cells; scale bar = 20 μm. Mast cells incubated with IgG from FMS patients exhibited a significant increase in surface-bound IgG, as demonstrated by fluorescence intensity compared to cells incubated with IgG from healthy controls. Colocalization of green (IgG) and red (mast cells) in PMCs was analysed using ImageJ’s Colocalization Finder, and the ICQ coefficient was measured. ****p < 0.0001 for FMS vs. HC. Data are expressed as mean ± SEM (n = 9/group for hind paws and n = 5/group for PMCs) and analysed using a student’s t-test. Created with BioRender.com.

### FMS-IgG has greater affinity for peritoneal mast cells relative to HC-IgG

To further investigate any systemic role of FMS-IgG on mast cells, peritoneal mast cells (PMCs) were collected on day 4 post-treatment and concentrated. Mast cells, human IgG and nuclei were stained using Immunofluorescence and Avidin-fluorescence and colocalised using ImageJ’s ‘colocalization finder’. PMCs treated with FMS-IgG exhibited increased surface-bound IgG compared to HC-IgG (Fig.2C-D).

The findings of mast cell accumulation in skin and of increased systemic FMS-IgG binding to mast cells indicate potential cell activation and therefore highlight that these cells might have relevance for the development of the FMS-model phenotype.

### Ablation of Mrgprb2-positive mast cells reduce hypersensitivity specifically in FMS-treated mice

To evaluate the impact of mast cell depletion on FMS-related pain behaviour, we again conducted behavioural tests using a von Frey filament assay to assess mechanical allodynia in mice treated with FMS-IgG. We utilised Cre-driven diphtheria toxin A (DTA) to selectively deplete Mrgprb2 (b2)-positive mast cells, specifically targeting all connective tissue mast cells, without affecting the circulating mucosal mast cells ^26, 27^. Mice treated with FMS-IgG following b2DTA-mediated mast cell depletion exhibited reduced (normalised) mechanical hypersensitivity by day 4 compared to wild type (WT) FMS-IgG-treated mice (Fig. 3A-B). Cold allodynia was also markedly reduced in b2DTA FMS mice (Fig. 3C). The absence of mast cells had no impact on faecal matter production (Fig. 3D), or on stool consistency measured with the Bristol stool chart (not shown). These data suggest that connective tissue mast cells are essential for the development of pain hypersensitivity in response to FMS-IgG, linking IgG induced mast cell activation directly to the pain symptoms in FMS. In contrast, IgG mediated abnormal stool production is not dependent on these cells.

**Figure 3.**
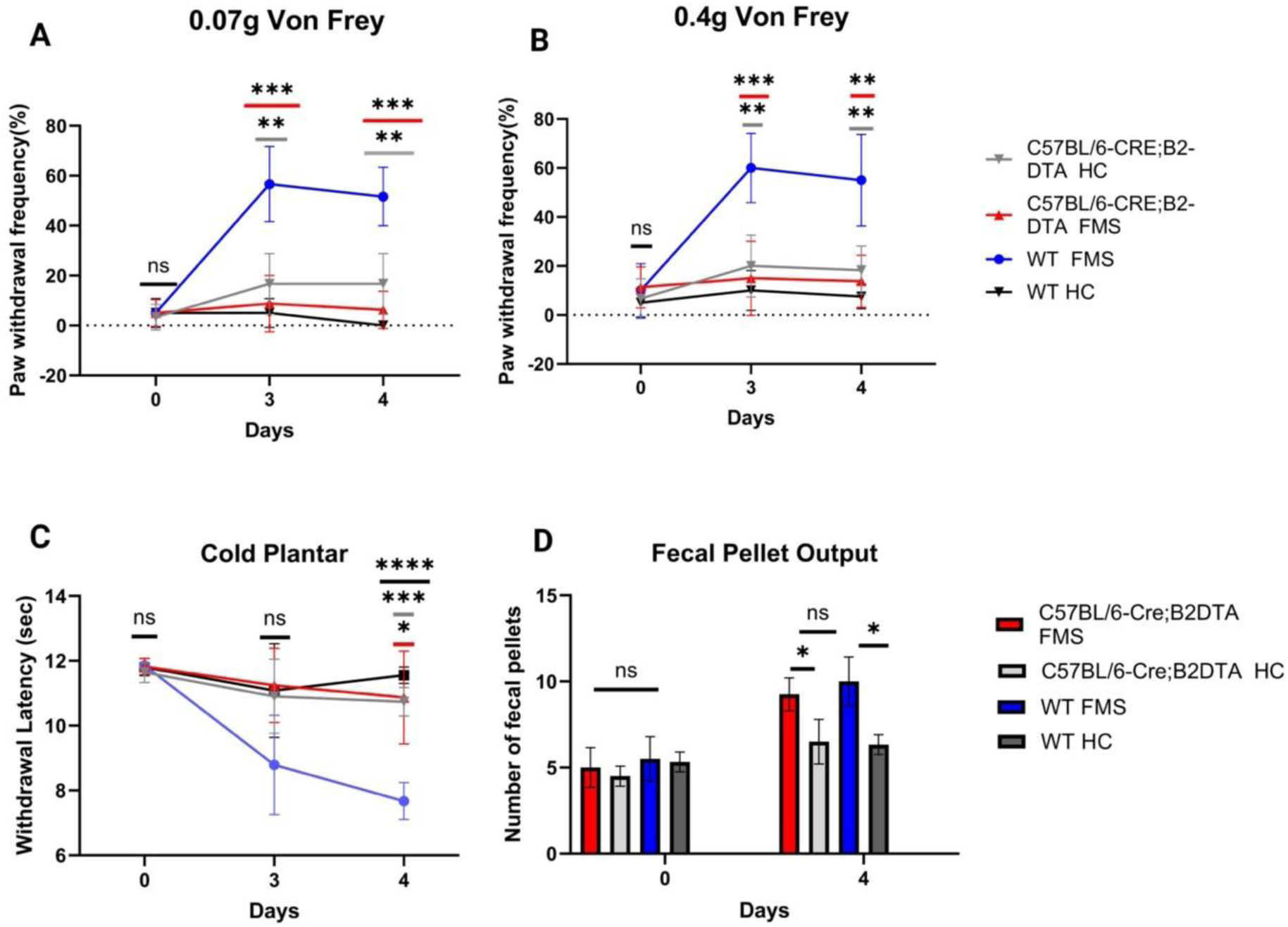
Ablation of Mrgprb2-positive mast cells reduced allodynia specifically in FMS-IgG treated mice. To confirm the role of mast cells in mediating the effects of FMS-IgG, we performed selective mast cell depletion in mice using Cre-driven diphtheria toxin A (DTA) to target Mrgprb2-positive mast cells. Human fibromyalgia IgG (8 mg) was administered via intraperitoneal injection (IP) for 2 consecutive days. Experimental design as in Figure 1A. (A-B) Mechanical sensitivity testing using Von Frey filaments on both hind paws showed a significant difference in response by day 4 between Mrgprb2DTA FMS mice and the other treatment groups. (C) Cold allodynia was assessed using the cold plantar assay on both hind paws, with a significant difference observed in Mrgprb2DTA FMS mice compared to the other treatments. (D) Stool production was quantified and compared across treatment groups. (n = 7 per treatment group for FMS). Statistical significance is indicated as ns = not significant, *p < 0.05, **p < 0.01, ***p < 0.001, and ****p < 0.0001 for WT FMS vs. WT HC vs. Mrgprb2DTA FMS vs. Mrgprb2DTA HC, analysed using one- or two-way ANOVA followed by Bonferroni’s correction. Data are presented as mean ± SEM. Created with BioRender.com.

### The absence of mast cells does not prevent pain development in CRPS-IgG-treated mice following injury

In persistent c<u>o</u>mplex Regional Pain Syndrome (CRPS), patients experience pain, allodynia, hyperalgesia, and persistent limb swelling long after the resolution of the inciting tissue injury. A CRPS IgG passive transfer-trauma model reproduces these features^26^. The experimental procedure is shown in (Fig. 4A). Mice treated with human CRPS IgG, after small hind paw trauma developed unilateral hind paw hyperalgesia to both mechanical and cold stimuli and increased hind paw swelling as previously described (Fig. 4 B-E); however, these mice did not exhibit any change in fecal output, unlike their FMS counterparts (Fig. 4 F-G). To evaluate the impact of mast cell depletion on CRPS-related pain behaviour, we repeated these tests using Cre-driven diphtheria toxin A (DTA) to selectively deplete b2-positive mast cells. The selective depletion of Mrgprb2-expressing mast cells produced no reduction in either mechanical or cold allodynia (Fig.4D) contrasting the findings with FMS transfer. Notably, a slight reduction of paw swelling in mast cell depleted mice was identified, potentially indicating diminished CRPS-related neurogenic inflammation (Fig.4 E). Mast cells do not appear to significantly influence the development or progression of CRPS-related hypersensitivities. These findings indicate a differential role of mast cells between CRPS and FMS.

**Figure 4.**
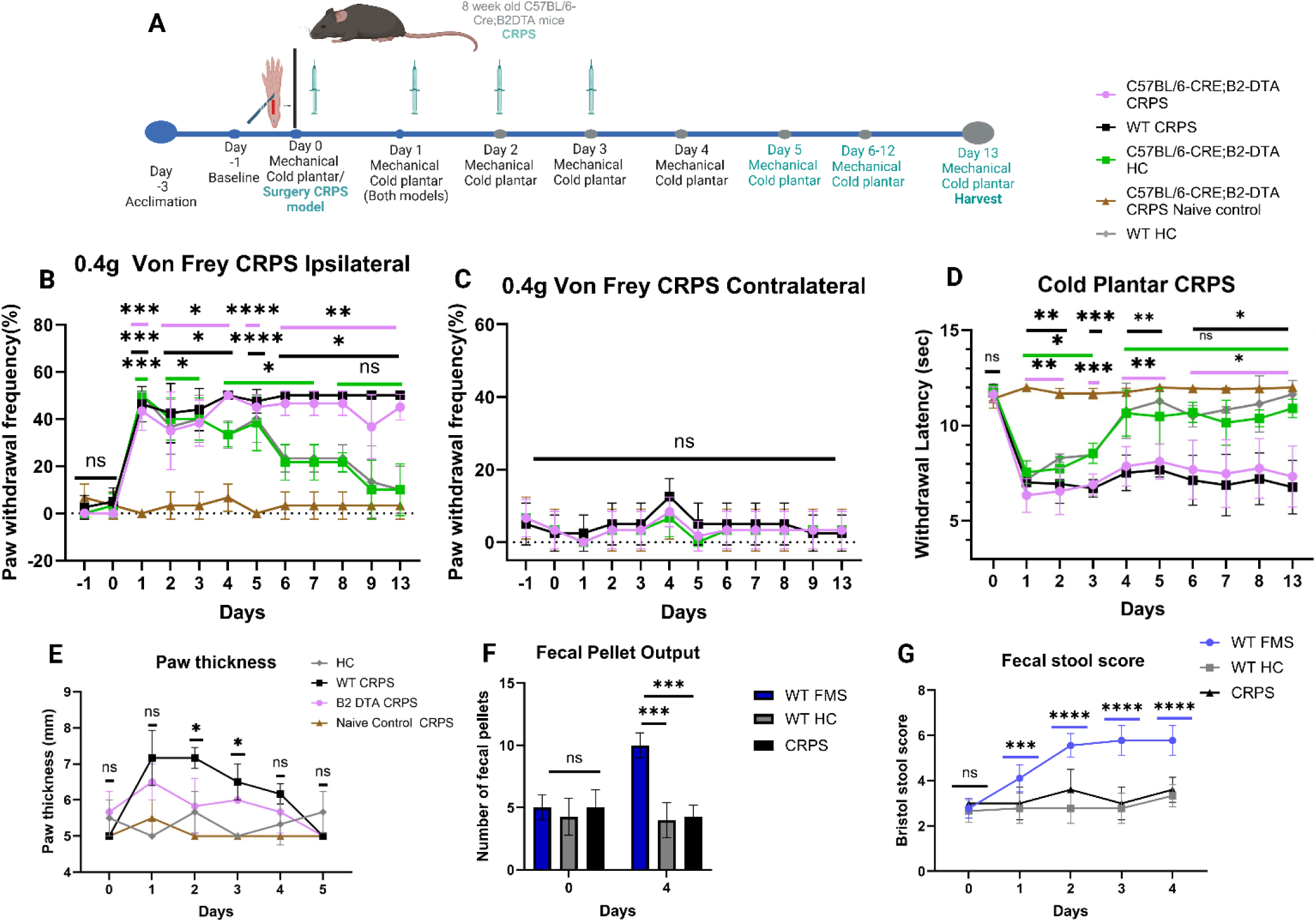
The absence of mast cells does not prevent pain development in CRPS-IgG-treated mice following injury. Mice were administered Human CRPS IgG via intraperitoneal injection (IP) 8mg for 4 consecutive days after plantar (skin-muscle) incision on the right paw. (A) Schematic illustration of the experimental design relating to the CRPS model. (B-C) Mechanical sensitivity testing using Von Frey filaments on both hind paws revealed a significant difference in response by day 4 between the Mrgprb2 DTA (Mrgprb2DTA) CRPS and WT CRPS groups and other treatment groups for the ipsilateral but not contralateral hind paws. (D) Cold allodynia was assessed using the cold plantar assay on both hind paws, with a significant difference observed between Mrgprb2DTA CRPS and WT CRPS and the other treatment groups. The data for behavioural nociception assays were compared to the naive control (no injury), and significance indicated by asterisks, with the colour bar corresponding to the treatment being compared. (E) Edema development in the hind paw was measured using callipers to assess paw thickness in millimetres. (F-G) Stool production was quantified and compared across different treatment groups. (n = 6 per treatment group for CRPS). Statistical significance is indicated as ns = not significant, *p < 0.05, **p < 0.01, ***p < 0.001, and ****p < 0.0001 for WT CRPS vs. WT HC vs. Mrgprb2 DTA CRPS vs. Mrgprb2 DTA HC, analysed using one- or two-way ANOVA followed by Bonferroni’s correction. Data are presented as mean ± SEM. Created with BioRender.com.

### Elimination of the mast cell receptor Mrgprb2 reduces hyperalgesia associated with FMS and normalises skin mast cell numbers

The Mrgprb2 receptor has previously been identified as a key player in the development and maintenance of several forms of inflammatory pain^27^. The role of this receptor in FMS-IgG mediated behaviour and mast cell activation was investigated using Mrgprb2 knock-out (b2KO) mice. We conducted mechanical paw withdrawal and cold plantar assays to assess mechanical and thermal allodynia in FMS-IgG treated mice. FMS-IgG treated b2KO mice exhibited reduced (normalised) mechanical and cold allodynia and reduced mast cell recruitment by day 4 relative to wild type (WT) FMS-IgG treated mice (Fig. 5A-E). These results highlight an essential role of the Mrgprb2 receptor for IgG-mediated pain sensitisation in this model.

**Figure 5.**
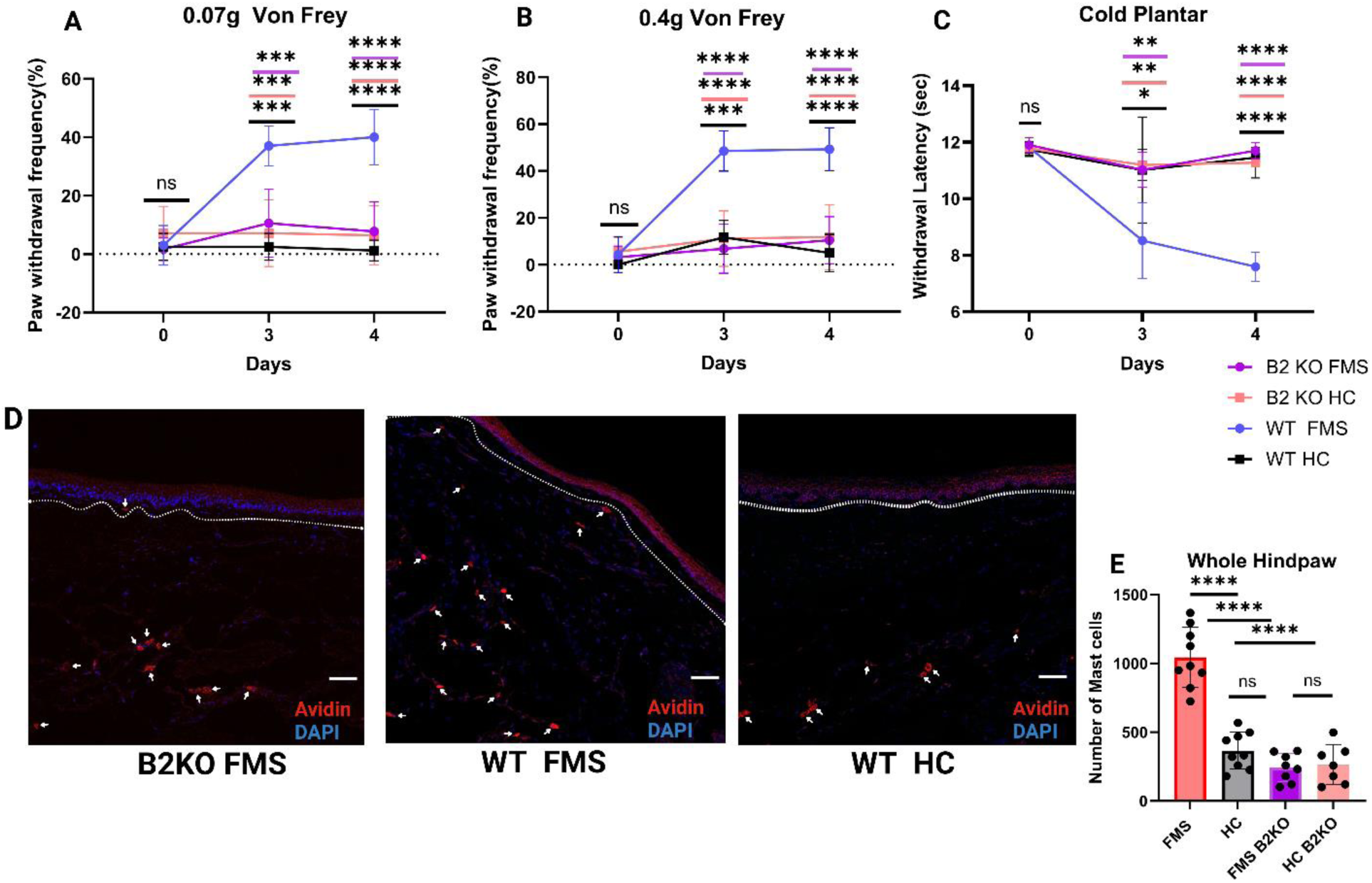
Elimination of the mast cell receptor Mrgprb2 reduces allodynia associated with FMS and normalises skin mast cell numbers. To validate the role of Mrgprb2 receptor on mast cells in mediating the effects of FMS-IgG, we deleted Mrgprb2 receptor. Experimental design as in Figure 1A. (A-B) Mechanical sensitivity testing with Von Frey filaments on both hind paws revealed a difference in response between b2KO mice and other treatment groups (pooled results from 2FMS and 2HC samples). (C) Cold allodynia was assessed using the cold plantar assay on both hind paws, with differences observed when comparing FMS treatment to other groups. (D) Immunofluorescence analysis of 14 μm sagittal sections of the hind paw was performed. Mast cells were stained in red (Avidin), and nuclei in blue (DAPI), scale bar = 100μm. Mast cell are indicated by white arrows, showing reduced mast cell recruitment in b2KO mice. (E) Quantification of mast cell numbers from immunofluorescence staining of hind paw sections. (n = 9 paws per treatment group for behaviour testing and staining). Statistical significance is indicated as ns = not significant, *p < 0.05, **p < 0.01, ***p < 0.001, and ****p < 0.0001 for WT FMS vs. WT HC vs. b2KO FMS vs. b2KO HC (colour coded), analysed using two-way ANOVA followed by Bonferroni’s correction. Data are presented as mean ± SEM. Created with BioRender.com.

### Quantification of FMS-IgG binding and activation of mast cell in the Presence or Absence of MRGPRX2

To investigate whether the activation of mast cells in the FMS-IgG transfer model may be caused by direct binding, we performed flow cytometry analysis on cultured human mast cells (LAD2), both WT and MRGPRX2-KO (KO) cells, after treatment with either FMS-IgG or HC-IgG. LAD2 mast cells incubated with FMS-IgG showed a higher proportion of IgG-positive cells (FcεRIα+ C-kit+) compared to those treated with HC-IgG (Fig. 6A). Notably, MRGPRX2 KO LAD2 cells demonstrated a significant reduction in FMS-IgG binding with all three tested FMS preparations (Fig. 6B-E). To understand whether IgG binding *in vitro* leads to mast cell activation, Elisa was performed to quantify secretion of IL-6, a proinflammatory cytokine released by mast cells which has previously been implicated in FMS^21^. The bind of FMS-IgG correlated with increased release of IL-6 in LAD2 wild-type (WT) compared to MRGPRX2-KO (knockout) (Fig. 6F). Overall, these findings suggest that IgG from FMS patients can activate MRGPRX2 receptors on mast cells to release IL-6, potentially contributing to a pro-nociceptive environment in FMS tissues. Further studies are needed to elucidate the mechanisms underlying this effect and the role of mast cell signalling pathways in FMS and other chronic pain conditions

**Figure 6.**
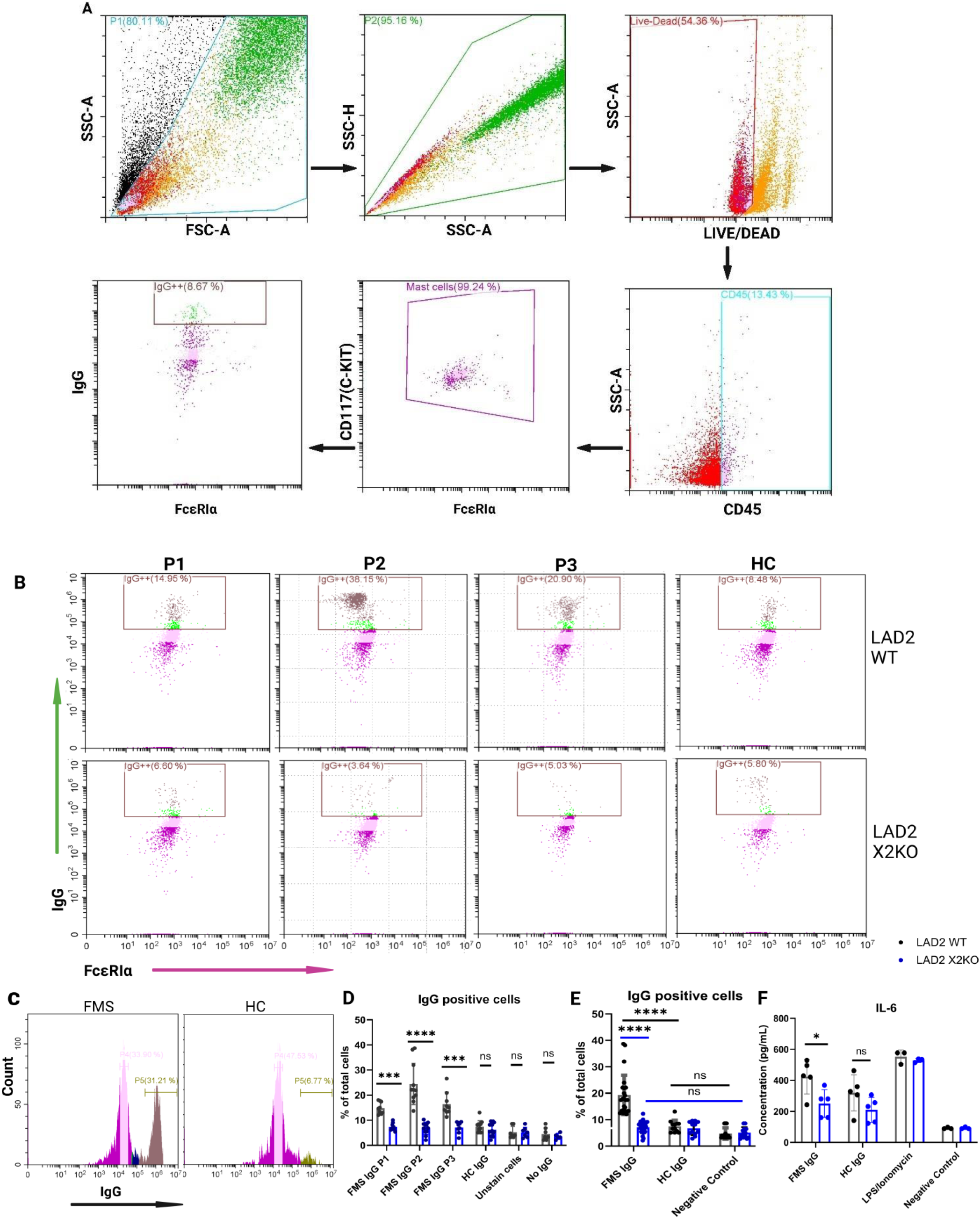
Quantification of Mast Cell Binding and Activation by FMS-IgG in the Presence or Absence of MRGPRX2. (A) Representative flow cytometry panels showing the percentage of mast cells pre-gated for FcεRIα and C-kit, used to identify mast cells and the population positive for IgG. Comparison made between wild-type LAD2 cells and LAD2 MRGPRX2 knock out mast cells (X2KO) after incubation with IgG from either FMS or HC patients. (B) Representative flow cytometry plots of LAD2 cells and LAD2X2 KO cells after treatment with IgG from various FMS patients (P1-P3), showing the population of mast cells (FcεRIα+C-Kit+) that are IgG positive (n=9 experiments/group). (C) Histogram representing the cell count of IgG binding in WT LAD2 mast cells. (D-E) Bar graphs showing the percentage of total cells that are IgG positive, comparing WT mast cells vs. X2KO mast cells, and FMS vs. HC. All data are presented as mean ± SEM of at least three independent experiments. (F) Bar graph showing the concentration of IL-6 released by LAD2 cells after IgG treatment comparing different groups (n=5/3 experiments/group). Comparisons between FMS patients and healthy controls, as well as between WT and X2KO mast cells, were analysed using one-way ANOVA followed by Dunnett’s post-hoc test. ns = not significant, *p < 0.05, ***p < 0.001, and ****p < 0.0001. Created with BioRender.com.

### Mast cell density is increased in FMS patient skin compared to healthy volunteers

Group differences in mast cell measures between patients with FMS and healthy volunteers (HV) were identified. Both the density and the total percentage of mast cells in biopsies from the proximal thigh were substantially higher in FMS patients relative to HV (Mast Cell Density: median [IQR]: FMS = 107.1 [66.7–138] cells/mm², HV = 46.4 [43.5–62.6] cells/mm²; p = 0.0006; Mast Cell Percentage: median [IQR]: FMS = 14.60% [9.1–18.6], HV = 9.3% [7.5–11.2]; p = 0.04) (Fig. 7A). Over half the participants with FMS (n=10) had mast cell densities greater than three standard deviations above the mean HV threshold (Fig.7B). Correlation analyses were performed between cutaneous mast cell densities and patients’ phenotypic FMS parameters as documented by the SFNSL and PDQ. The patients’ skin mast cell densities showed significant correlations of mild to moderate strength with several of these parameters, whereas inverse correlations were rarely observed (Table S1).

**Figure 7.**
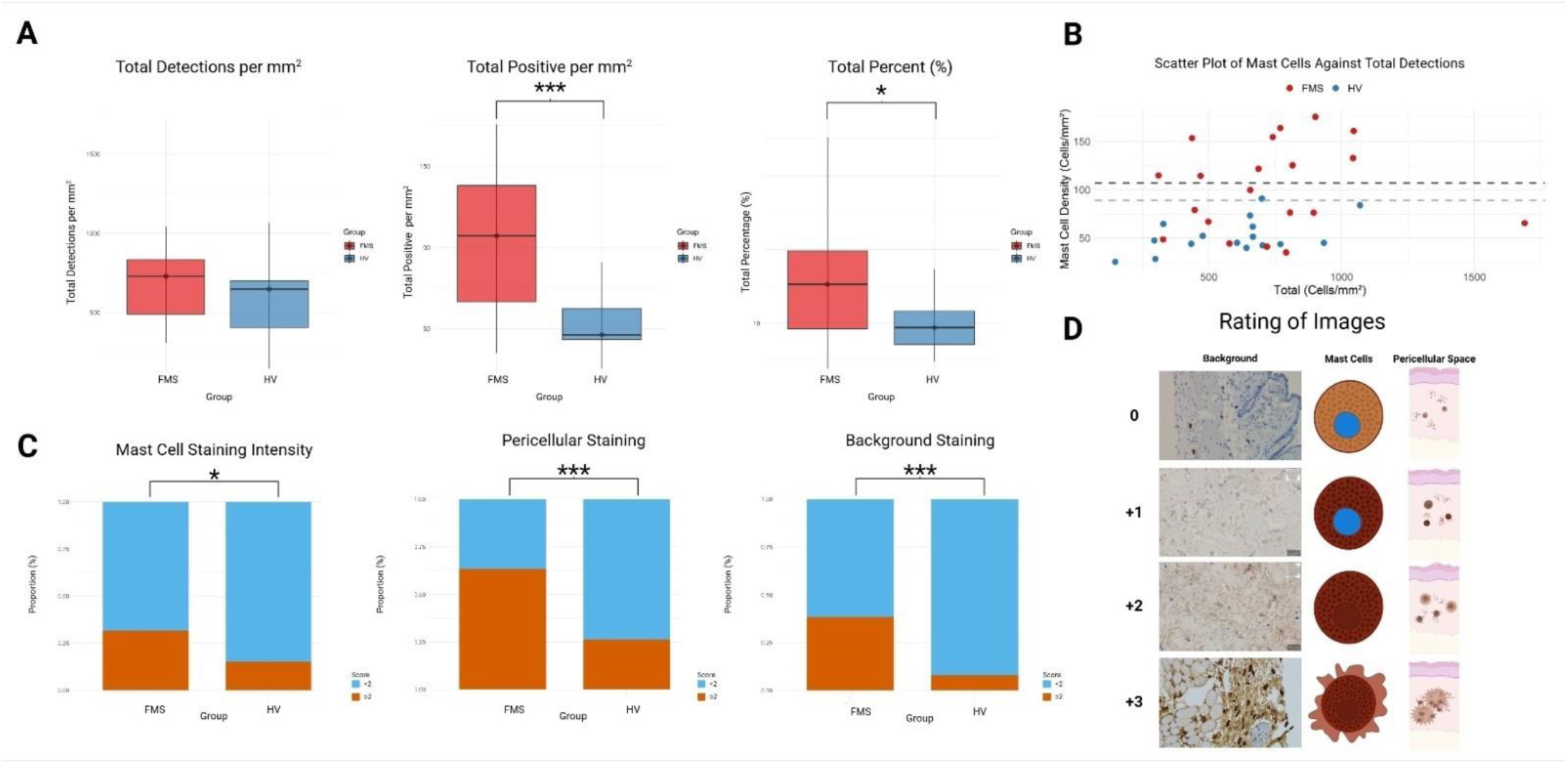
Mast Cell Density is Abnormal in Human FMS Skin. (A) Box plots comparing total number of cells (Left), mast cell densities (Middle) and mast cell percentage (Right) between two groups: FMS (red) and HV (blue). The median is represented on the centre line with the error bars indicating the inter-quartile range at 0.25 to 0.75. Participants with FMS had significantly increased cell densities and a higher positive percentage of mastocytes relative to HV. (B) Scatter plot of mast cell densities against total number of cells within tissue sections for participants with FMS and HV. About half of participants with FMS (n=11) had mast cell densities greater than three standard deviations above the mean HV threshold. The red dots represent participants with FMS whilst the blue dots represent HV participants. The red dashed line represents the 2 standard deviations (SD) threshold for HV while the blue dashed line represents the 3 SD threshold for HV. (C) Bar plots comparing Mast Cell Staining Intensity (Left), Pericellular Staining Intensity (Middle), and Background Staining (Right) between HV (left bars) and FMS (right bars) groups. Blue bars represent scores less than 2, while orange bars indicate scores of 2 or higher. (D) *The grading of mast cell staining outcomes across four levels* (0, +1, +2, +3), based on three key parameters: background staining, mast cell staining intensity, and pericellular space staining. These parameters are graded independently, allowing for different combinations across a single examined image. *Grade 0:* Minimal background staining, mast cell staining with the nuclei clearly visible, no pericellular staining. *Grade +1:* Minimal background staining, darker mast cell staining with the nuclei visible, and slight staining of the pericellular space. *Grade +2:* Noticeable background staining, intense mast cell staining, nuclei is obscured by chromogen, pronounced pericellular space staining. *Grade +3:* More prominent background, dark mast cell staining, increased, staining in the pericellular space. Statistical significance is indicated as follows: *p < 0.05, **p < 0.01, ***p < 0.001. Created with BioRender.com.

### Anti-tryptase staining is enhanced in FMS skin relative to HV

The AA1 antibody used to stain for mast cells in human skin unexpectedly often also stained tissue around the mast cells. The FMS group showed a significantly higher proportion of elevated assessment scores (≥2) compared to the control group for mast cell staining intensity (FMS = 31.7% vs. HV = 14.9%; +16.7%, p = 0.027), background staining (FMS = 38.3% vs. HV = 8.0%; +30.3%, p < 0.0001), and pericellular staining (FMS = 63.3% vs. HV = 26.4%; +36.9%, p < 0.0001) (Fig 7C). These findings suggest that tryptase isoforms are elevated in FMS skin, likely released from mast cells, and may be an important biomarker in FMS.

## DISCUSSION

Previous work has demonstrated that patients with severe fibromyalgia syndrome (FMS) invariably harbour pathogenic serum IgG autoantibodies which, upon passive transfer to mice elicit core elements of the condition including mechanical and cold hypersensitivities, reduced grip strength and locomotion, and small skin fibre pathology. Increased FMS-IgG binding to satellite glia cells was demonstrated^6, 28, 29^. However, direct evidence relating to mechanisms through which pathogenic IgG exert their effects has been lacking. The present study provides compelling evidence that the pathogenic effect of FMS-IgG on mechanical and cold hypersensitivities is mediated by mast cells and that a crucial role is played by the MRGPRX2 receptor.

We first provided evidence from behavioural studies which validates previous findings on mechanical and cold hypersensitivities, both through reproduction and by demonstrating via *in vivo* imaging of dorsal root ganglion cells that IgG-induced sensitisation of nociceptive afferents is reflected in increased responsiveness of neuronal DRG cell bodies *in vivo*. Increased skin mast cell density in FMS has historically been reported by two independent groups^13, 14^; mast cell accumulation may indicate activation of these cells^9, 30^. We hypothesised that pathogenic FMS-IgG may cause skin mast cell accumulation, and further, that mast cells may cause pertinent elements of the IgG transfer-induced FMS phenotype.

To investigate this, we first conducted immunofluorescence and immunochemistry experiments showing that mast cell numbers in the skin of FMS-IgG transfer mice are indeed significantly increased; our results are consistent with the historical results in patient skin and with our hypothesis. Increased mast cell density was identified in both glabrous and non-glabrous skin indicating widespread IgG effects. We next demonstrated increased binding of FMS-IgG to peritoneal mast cells *in vivo* suggesting that direct IgG binding may contribute to abnormal mast cell recruitment. When we subsequently probed the effect of connective tissue mast cell knock out using Mrgprb2DTA mice we found that the sensitising effects of FMS patient IgG are abrogated suggesting that mast cells are essential to their development. These changes were consistently observed after injection of preparations from each of three different FMS donors.

However, we also discovered that mast cell involvement is not ubiquitous in autoantibody mediated chronic pain. Complex Regional Pain Syndrome is a severe chronic limb pain condition typically triggered by tissue injury or trauma to a distal limb and associated with localised increased neuropeptide release and neuroinflammation ^31^. Clinically, CRPS is considered a ‘sister’ condition to Fibromyalgia, with both conditions categorised under the umbrella ‘primary chronic pain’ in the WHO ICD-11 classification system. These are conditions with pain as the primary clinical problem, rather than pain caused by other disease or tissue change^32^. Prior research demonstrated that persistent CRPS may also be caused by pathogenic serum-IgG, but which become harmful only in the context of limb injury, a key difference from FMS-IgG^33–35^. We now show that the skin mast cell density is normal in this CRPS ‘passive-transfer-trauma model’, and the pain phenotype is unaffected by mast cell knockout. This is consistent with earlier, independent investigations indicating that the skin mast cell density in the affected skin of *patients* with *persistent* CRPS is normal, highlighting probable lack of abnormal activation, contrasting the findings in FMS skin^36, 37^. Mast cells appear to be a key player in the pathogenesis of FMS, but not necessarily in other chronic pain conditions even when they are associated with neuroinflammation.

Activation of the MRGPRX2 or Mrgprb2 receptor on mast cells triggers a predominantly IgE-*independent* activation pathway, leading to degranulation. This pathway plays a key role in conditions where mast cell activation occurs unrelated to allergens. It results in the release of substances like histamine, proteases, cytokines (IL-6), and chemokines that contribute to inflammation and pain^17^. The release of substance P from peptidergic nociceptors has previously been demonstrated to result in the recruitment of mast cells, in addition to activating the MRGPRX2/b2 receptor^9, 21, 22^. With this in mind we probed the effect of selective Mrgprb2 knockout and found that the FMS phenotype was abrogated in Mrgprb2—FMS-IgG injected animals. The skin mast cell density was normalised, strongly suggesting that Mrgprb2 receptors are involved in mediating the pathogenic effects of FMS-IgG on mast cells.

The MRGPRX2/b2 receptor has not previously been identified as critical for mast cell activation in immune responses mediated by IgG, to our knowledge. Such mast cell activation might result from direct receptor binding or, alternatively the receptor may be part of a signalling pathway complex that facilitates downstream effects of FMS-IgG binding to other receptors like FcγRs ^20^. Therefore, investigating any direct binding and activation of FMS-IgG autoantibodies to mast cells in an MRGPRX2-dependent manner was crucial for further clarification of the mechanism of its involvement in causing hypersensitivities. To investigate the role of the MRGPRX2/b2 receptor in IgG binding, we utilized the human LAD2 cell line. We found that wildtype LAD2 cells treated with FMS-IgG exhibited stronger IgG binding compared to those treated with HC-IgG. Furthermore, knocking out MRGPRX2 in LAD2 (X2-KO) cells reduced the IgG positive population. To understand whether IgG binding leads to activation of MRGPRX2 in mast cells *in vitro*, ELISA was performed to detect IL-6. An increase in IL-6 secretion in WT cells treated with IgG from FMS patients compared to X2-KO cells was observed. This supports the idea that MRGPRX2/b2 may play a role in mast cell-mediated IL-6 release in FMS patients^21^. Interleukin-6 (IL-6) is a cytokine that is crucial in the role of pain modulation and the regulation of immune cell activity, including mast cell proliferation^38, 39^. Thus, the interplay between IL-6 and mast provides an explanation for the increase of mast cell observe in FMS patients and FMS treated mice.

Our findings demonstrate for the first time that connective tissue mast cells are a critical element facilitating FMS IgG transfer-mediated hypersensitivities to the core FMS phenotype elements of mechanical and cold stimuli. We further provide evidence that CTMCs are directly bound by FMS-IgG and that this process requires the presence of the Mrgprb2 receptor. The presence of MRGPRX2/b2 on cells also leads to raised secretion of IL-6, a key pro-inflammatory cytokine associated with autoimmune disease with potential for therapeutic options ^21^. One limitation to the interpretation of IgG-induced IL-6 secretion in LAD2 cell is that various other receptors like Fc gamma receptors (FcγRs) can also cause release of IL-6. Therefore, we cannot rule out the possibility that the MRGPRX2 receptor may function in coordination with other receptors. FcγRs have been linked to IgG binding in autoimmune diseases^40, 41^. Studies showing the synergistic interaction between MRGPRX2 and FcεRI via IL-33 suggest that FcγRs may also contribute to this mechanism^42^.

Having established an important role for mast cells upon FMS-IgG transfer, we have then forward translated our experimental findings by examining thigh skin of a UK cohort of FMS patients and controls. We were able to both confirm historical findings of strongly increased skin mast cell density in FMS skin for the first time in UK patients, and to provide preliminary evidence of a correlation with pertinent FMS disease parameters including pain intensity, pressure and temperature sensitivity, urinary- and GI-symptoms (Table S1). We further show, for the first time evidence for another mast cell activation marker in patient skin, increased intra- and extra-cellular tryptase.

Overall, our findings indicate that FMS-IgG has unique properties that promote the activation of connective tissue mast cells (CTMC), a process which requires the presence of the MRGPRX2 receptor. This activation may drive changes critical to the IgG transfer phenotype and potentially play a key role in the clinical manifestation of FMS. These results highlight the ability of FMS-IgG to differentially activate mast cells, leading to IL-6 release and potentially contributing to a pro-nociceptive tissue environment in FMS. The MRGPRX2 receptor appears to be a promising therapeutic target for FMS; blocking or modulating its function could potentially alleviate symptoms experienced by patients suffering from FMS.

## METHODS

This study was designed to investigate the mechanism by which FMS serum-IgG leads to pathogenic effects via the activation of connective tissue mast cells (CTMC) and the mast cell MRGPRX2/b2 receptor. We utilized a combination of microscopy-based imaging, cell analysis and behavioral approaches both *in vivo* and *in vitro,* utilizing both animal models and human samples. IgG isolation from multiple individuals with fibromyalgia syndrome (FMS), healthy controls (HC) and one complex regional pain syndrome (CRPS) patient was used to assess mast cell activation, binding, and the development of allodynia. All experiments were performed in replicated per condition, and data was analyzed using various statistical methods appropriate for the sample size.

### Patient IgG donors and IgG purification

Plasma for the passive transfer experiments was derived from waste plasma available from three FMS patients and one CRPS patient who harboured pathogenic IgG autoantibodies as demonstrated in earlier passive-transfer experiments^6, 34^, and who had received clinical plasma exchange to treat their pain condition. Patients individually consented to the retainment and storage of their waste plasma. University ethical approval to utilise waste plasma for the current studies was obtained (UoL 12267).

The patients were three females, in their thirties, forties (CRPS), and fifties respectively, and one male in his forties; all patients had been diagnosed >5 years before plasma donation. The FMS patients fulfilled FMS ACR 2016 criteria and the CRPS patient met Budapest research criteria^43, 44^. The patients’ average (worst) daily pain intensities ranged between numeric rating scale (NRS) 6(8), an FMS patient – 8.2(9.5), the CRPS patient. Plasma IgG was purified using Protein G large columns, and the IgG concentration was adjusted to 20mg/ml in normal saline as previously described^6, 33^. Purified IgG was stored at 4C and shipped on ice for in vivo experiments within 2 weeks.

### Mice strains

We transferred patient IgG from either of three patients (two females) with severe FMS, one patient with severe persistent CRPS, or age/sex-matched healthy controls to groups of 2–4-month-old male and female C57BL/6 background mice. Pirt-Cre; GCaMP6s mice were obtained by crossing Pirt-Cre mice with Rosa26-lox-stop-lox GCaMP6s animals^45^. Mrgprb2 DTA mice were generated by Mrgprb2Cre transgenic mice crossed with Cre-dependent ROSA26dta mice. Mrgprb2 KO was generated using CRISPR/cas9 targeting Mrgprb2^46^. WT littermates were also used in behavioural experiments along with generated strain mice. All experiments were performed with the protocols approved by the Animal Care and Use Committee of Johns Hopkins University School of Medicine. The mice were housed in the vivarium with a 12-h light/dark cycle and all mice were acclimated for at least 30 min to their testing environment. Mice were housed 4-5 mice in each cage in the vivarium.

### Immunoglobulin treatment

Mice were injected intraperitoneal (i.p) with 8 mg/mL IgG from HC subjects or FMS patients on 2 consecutive days. CRPS treatment: Mice were injected intraperitoneally (i.p) with 8 mg/mL IgG from HC subjects or CRPS patients on 5 consecutive days after classical plantar incision^47^ on the right paw. This procedure (passive transfer followed by hind paw incision) was previously shown to cause persistent isolated right hind paw hyperalgesia and swelling^34^. Paw thickness/swelling in the CRPS model was measured daily after injection using caliper^48^.

### Withdrawal threshold to mechanical stimulation

To measure mechanical sensitivity, each mouse was placed in a Plexiglas chamber on an elevated mesh screen and was assessed with the von Frey test by the frequency method. Two calibrated von Frey monofilaments (low force = 0.07 g; high force = 0.4 g) were used. Each von Frey filament was applied perpendicularly to the plantar side of each hind paw for ∼1 s until the fibre bent. The stimulation was repeated 10 times every 1 second with at least 5 minutes pause prior to testing of the other paw of the same animal. A positive response was defined as the rapid withdrawal of the paw from the filament. The occurrence of withdrawal in these 10 trials was expressed as a percent response frequency: paw withdrawal frequency (PWF) = (number of paw withdrawals/10 trials) × 100^49^.

### Cold sensitivity

The cold plantar assay was used to evaluate noxious cold sensitivity. Mice were placed individually into acrylic containers separated by opaque dividers on a 3/16″ glass plate and allowed to acclimate for 30 minutes before being tested. Powdered dry ice in a cutoff 10-mL syringe was held against the glass beneath the hind paw until the paw was withdrawn. Applications were repeated at 5-minute intervals, alternating paws, for a total of 4 trials. The mean paw withdrawal latency (PWL) was calculated from these replicate measurements, the data was averaged for each group as described^50^. To avoid potential tissue damage, the cutoff maximum stimulus time was 20 sec for mice in most cases.

### In vivo DRG calcium imaging

In vivo imaging of whole L4 DRG in live mice was performed for 2–7 hr immediately after the exposure surgery (as previously described^45^). Body temperature was maintained at 37°C ± 0.5°C on a heating pad and rectal temperature was monitored. After exposure surgery, mice were laid down in the abdomen-down position on a custom-designed microscope stage. The spinal column was stabilised using purpose-built clamps to minimise movements caused by respiration and cardiac activity. In addition, a specially designed head holder was also used as an anaesthesia/gas mask. The animals were maintained under continuous anaesthesia for the duration of the imaging experiment with 1%–2% isoflurane gas using a gas vaporiser. Pure oxygen was used to deliver the gas to the animal ^45^.

The microscope stage was fixed under a laser-scanning confocal microscope (Leica LSI microscope system). Live images were acquired at typically ten frames with 600 Hz in frame-scan mode per 7 s with solid diode lasers (Leica) tuned at excitation 488 nm wavelength and emission at 500–549 nm for green fluorescence raw imaging frame and were imported into ImageJ (NIH) for further analysis. DRG neurons were at the focal plane, and imaging was monitored during the spontaneous activation of DRG neuron cell bodies without peripheral stimuli. The imaging parameters were chosen to allow repeated imaging of the same cell for a long time without causing damage to the imaged cells or surrounding tissue. At the end of each experiment, the viability of the DRG was confirmed by observing large numbers of cells giving a calcium transient in response to pinching of the hind paw. GCaMP6s was chosen as a highly sensitive calcium indicator with slow kinetics as it can detect single action potentials in vivo under optimised recording conditions. We recorded the activated neurons by brushing and pinching the hind paw before (1st) and after (2nd) application of 1.5% lidocaine onto the DRG. We found that lidocaine can inhibit all neurons’ activation by brushing and pinching. 3 hours after washing out lidocaine, we brushed and pinched again (3rd), when we found that the neurons had recovered.

### Stool collection and analysis

The paper towel method was used to collect mice stools. Mice were placed on a wire screen and separated by acrylic opaque dividers for 30 minutes. To eliminate the influence of the biological rhythms, the assay was conducted at 8:00 am once daily. After collection, the changes in stool form (pellet size and consistency) were observed and recorded^45^. We used the Bristol Stool Chart to assess variations in water content in faecal matter, allowing us to identify potential gastrointestinal changes based on stool consistency.

### Staining of mice paws

Adult mice were anesthetised and perfused with 30 mL 0.1 M phosphate buffered saline (PBS), 4°C followed with 30 mL of fixative 4% paraformaldehyde, 4°C. Hind paw skin tissue was dissected removing the hypodermis, ligaments and muscle tissue and stretched out using insect pins. The samples were then fixed at room temperature (RT) for 45-60 minutes in 4% buffered paraformaldehyde (PFA). Tissues were cryoprotected in 20% sucrose for up to 8 h followed by 30% sucrose for 24 h and then sectioned (14 µm width) with a cryostat. The slides were pre-incubated in blocking solution (10% normal goat serum (vol/vol), 0.03% Triton X-100 (vol/vol) in PBS, pH 7.4) for 1h at RT. The staining included the use of primary antibodies against anti-Human IgG conjugated, Alexa Fluor 488 (Invitrogen,1:400 dilution, A-11013), Mast Cell Tryptase Antibody (AA1) Alexa Fluor® 594 (Santa Cruz,1:500 dilution, sc-59587) or Avidin-Sulforhodamine 101 (Avidin-Texas Red) (Sigma Aldrich, 1:250 dilution, A2348), and Claudin 1 Monoclonal Antibody conjugated, Alexa Fluor 488 (Invitrogen,1:500 dilution, 374988). Sections were washed three times with PBS and Fluoromount G with DAPI (Invitrogen, 00-4959-52) was applied before coverslips were placed over section.

### Staining of non-glabrous skin

Adult mice (n=1 each) were injected for three consecutive days intraperitoneally with daily 8 mg of one of the three protein-G affinity-purified HC-IgG or FMS-IgG preparations respectively that had been used for the behavioural assays. On day 4, after humane killing non-glabrous skin tissue from the back of each mouse was collected and fixed at for 48 hours in 4% PFA. Specimens were processed to paraffin wax and microtomy was performed at 4µm. The tissue sections were stained with anti-tryptase 1 (AA1) using Anti-Tryptase antibody [EPR9522] (Abcam, 1:200 dilution; 1.145 µg/mL; ab151757), following standardised low-pH antigen retrieval, blocking and immunohistochemistry protocol visualised with 3,3′-Diaminobenzidine (DAB).

### Staining of Peritoneal mast cells

Animals were anesthetised via inhalation followed by cervical dislocation. Peritoneal mast cells were isolated by adding PBS to peritoneal cavity, platted into the well of a six-well plate, cells were allowed to recover for 2 h in DMEM at 37°C and 5% CO2. Cells were then spun at 1000 rpm for 5 min at 4°C on a Cytospin (Thermo Scientific), fixed with Cyto-Fast™ Fix/Perm Buffer Set (BioLegend, 426803) at RT for 20 min, stained with 1:250 Avidin-Texas Red and anti-Human IgG conjugated, Alexa Fluor 488 for overnight (12hrs) at 4°C, and were washed three times with PBS and Fluoromount G with DAPI (Invitrogen, 00-4959-52) was applied before coverslips were placed before imaging.

### Human mast cell culture

LAD2 (Laboratory of Allergic Diseases 2) been shown to express MRGPRX2 endogenously and LAD2 X2-KO (Laboratory of Allergic Diseases 2-MRGPRX2 KO)^51^ cells human mast cells were cultured in StemPro-34 SFM medium (Life Technologies) supplemented with 2 mM L-glutamine (Fisher), 100 U/mL penicillin (Fisher), 50 µg per mL of streptomycin (Fisher) and 100 ng/mL recombinant human stem cell factor (Peprotech). The cell suspensions were seeded at a density of a million cells per mL and maintained at 37 °C and 5% CO2.

### Flow cytometry

LAD2 or LAD2 X2-KO cells were collected and spun at 200 g for 5 min at 4°C. The cell pellets were resuspended in 1 mL PBS the cells were treated with IgG 10ug/mL then fixed using cyto-fast Fix/Perm buffer (BioLegend). Cells were then stained with 2 μL Live/Dead Fixable Aqua Dead cell stain in 1 mL of PBS for 30 min on ice. The samples were centrifuged at 200 g for 5 min at 4°C, resuspended in 100 μL ice-cold FACs buffer (1X PBS-2% FBS) and then Fc blocked for 10 min on ice. The cells were stained for 30 min with indicated antibodies, spun at 200 g for 5 min at 4°C and washed twice with 1 mL ice-cold FACs buffer. Final cell pellets were suspended in 300 μL FACs buffer, run on cytoFLEX (Beckman Coulter) and analysed using FlowJo v.10 software.

The following antibodies from Biolegend were used in flow analysis: goat anti-human CD45 (clone HI30, BioLegend) PE, goat anti-human CD117 (c-Kit, Biolegend) PE/Dazzle, goat anti-human FcεRIα (AER-37-BioLegend) Apc/Cy7, goat anti-human IgG (A11013, Thermo fisher scientific) The cells were initially gated for single cells based on forward and side scatter (FSC-A/SSC-A) followed by two double exclusion gates (SSC-A/SSC-H and FSC-H/FSC-W). Then, dead cells were excluded based on their positivity for Live/Dead Fixable Aqua Dead cell staining. Live cells were then gated for CD45+ cells. Mast cells were identified as CD117+FcεRI+.

### ELISA (enzyme-linked immunosorbent assay)

Cytokine concentrations from cell-free supernatants was quantified using commercially available ELISA kits specific for human IL-6 (DY206, R&D Systems). The detection limit for IL-6 was 600 pg/mL. LAD2 or LAD2 X2-KO cells were cultured for 24 hours before treatment with either FMS, or HC IgG (10 μg/mL) with a positive control combination of ionomycin (1 μM) and LPS (1 μg/mL) cultured for 6 hours.

### Detection of Mast Cells in FMS skin

#### Patients

Participants with Fibromyalgia Syndrome (FMS), and healthy volunteers (HV) were recruited. A priori ethical approval was obtained (Southwest - Frenchay Research Ethics Committee REC reference: 20/SW/0138). All participants provided written informed consent. Details are described elsewhere^52, 53^.

#### Questionnaires

Symptom burden and pain characteristics were assessed in all participants using the short-form McGill Pain Questionnaire ^53, 54^ (MPQ-SF), PainDetect^55^ (PDQ), Small Fibre Neuropathy Screening List^56, 57^ (SFNSL) and the revised Fibromyalgia Impact Questionnaire^58^ (FIQR).

### Quantitative Sensory Testing

All participants underwent quantitative sensory testing (QST) of the right forearm according to the German Research Network on Neuropathic Pain (DNFS) protocol^59^. Collected QST data was log and z-transformed to standardise all values with the exception of paradoxical heat sensations and dynamic mechanical allodynia as previously described^60^. This is a standard research methodology allowing comparisons between participants differing in age and sex^61^.

### Skin Biopsy Collection and Processing

Participants with FMS (n=20) and HV (n=16) underwent punch biopsies of the thigh. Specimens were sampled and handled as previously described^52^. Skin biopsies were subsequently processed and embedded into paraffin wax as shown in Fig. S3.

### Cutaneous mast cell quantification

Slides from patients with FMS (n = 20) and healthy volunteers (HV) (n = 16) were processed using a standardised automated immunohistochemistry protocol (Leica Biosystems, IL, USA, Bond RXm Autostainer) using ready-to use Anti-tryptase 1 (AA1) antibody (Agilent Technologies, Santa Clara, CA, USA) and were visualised with a DAB detection kit (Leica Biosystems, IL, USA, Bond Polymer Refine Detection kit DS9800). All tissue sections were imaged using an automated slide scanner (Zeiss Axioscan Z1, Carl Zeiss Ltd., Cambridge, UK) with a 20x objective (Plan-Apochromat 20x/NA 0.8) and quantified within 4µm paraffin-embedded tissue sections. Mast cell density was determined using the open-source software QuPath (Version 0.5.1) by applying a standardised protocol for cell detection, which used a single-gated threshold to classify positive and negative cells stained with haematoxylin and AA1 using DAB^61, 62^. To assess differences in staining intensity between groups, two histology-naïve raters, and one expert examiner, blinded to group assignments, evaluated the images based on background staining, mast cell staining, and peri-cellular space staining intensity, each rated on a Likert scale (0–3). The criteria used for scoring are visualised in Fig. 7D.

### Statistical analysis

#### Statistical Analysis: Immunofluorescent Rodent and Cellular Experiments

Data were analysed using GraphPad Prism (v7.0d) and presented descriptive statistics as mean ± standard error of the mean (SEM) where n represents the number of mice in the analysis. The distribution of all variables was assumed to be normal. Inferential group comparisons were either two-tailed, unpaired Student’s t-test (2 groups) or one- or two-way ANOVA with Bonferroni correction (> 2 groups). Statistical significance was defined as p < 0.05.

A statistically significant difference was defined as p < 0.05. Error bars on all plots are SEM, with statistical significance defined as *p<0.05,* ∗∗p<0.001, ∗∗∗∗p<0.0001.

#### Statistical Analysis: Brightfield Mast Cell Quantification

Data were analysed using RStudio (version 4.0.2) for the analysis of scanned tissue sections. The total number of detections of any cell nuclei per mm² (detections/area), total AA1-positive mast cells per mm² (mast cells/area) and the cumulative mast cell percentage (percentage of mast cells relative to total detections). Group comparisons were conducted using the Mann-Whitney U test for continuous variables for the human skin samples. For the rodent skin an analysis of variance (ANOVA) was used to perform group comparisons of mast cell density. Post-hoc pairwise comparisons were performed using Tukey’s Honest Significant Difference (HSD) test to identify specific group differences and control for multiple comparisons. Inter-rater reliability between the three raters was calculated using a two-way mixed-effects model which used the ‘irr’ package^63^. Delta values were calculated using the expert examiner as the reference. Once reliability was confirmed, the scores were dichotomised into <2 and ≥2 rating categories and summarised as frequencies and percentages for comparison using Chi-Square tests. Differences between raters were evaluated using the Wilcoxon signed-rank test. Pearson and Spearman correlation analyses were performed to assess the relationship between skin mast cell density and phenotypic FMS parameters effect sizes expressed as correlation coefficients (R) and determination values (R^2^)

## Funding

Howard Hughes Medical Institute (XD) Pain Relief Foundation, Liverpool (AG) Versus Arthritis Grant number 22471 (UA)

## Authors Contributions

Conceptualization: XD, AG

Methodology: KRS, JB, QZ, UA, HN, RB, DA, SK, AM

Investigation: JB, KRS, HN

Visualization: KRS, JB, QZ, UA, HN, RB, DA, SK, AM

Funding acquisition: AG, XD

Project administration: HN, DA, SK

Supervision: XD, AG

Writing – original draft: KRS, JB, HN, RB, DA, SK, AM, AG, XD

Writing – review & editing: JB, KRS, DA, RB, AU XD, AG

Correspondence to Xinzhong Dong for all animal experimental data, and to Andreas Goebel for all human data and rodent mast cell bright field assessment.

## Animal Study

Study conception and design X.D and K.R.S. Study funding was obtained by X.D. Animal behavioural experiments, histology and flow cytometry were conducted by K.R.S.. K.R.S performed analysis of all experimental data with the exception of mast cell bright field assessment conducted by JB with support from HN and SK.

## Human Study

Human waste plasma was provided by AG for the purpose of this study from material derived from his patients’ clinical treatment. All IgG purification was conducted by HN. Funding and study design for the DEFINE-FMS study was obtained by U.A. subsequent funding and supervision of immunohistochemistry was provided by A.G. A.M collected all questionnaire and quantitative sensory testing data. Histology and immunohistochemistry assays of human tissues was designed and conducted and optimised by J.B. Data analysis was undertaken by J.B.

## Competing interests

X.D. is the scientific founder of and consultant for Escient Pharmaceuticals, a pharmaceutical company developing drugs targeting Mrgprs. X.D. collaborates with GlaxoSmithKline (GSK) on Mrgpr-related projects unrelated to this manuscript. Other authors declare no competing interests.

## Data availability

The data that support the findings of this study are available from the corresponding authors upon reasonable request. All figures created using BioRender.com. Raw clinical data is protected and limited release may require data transfer agreement and associated costs.

## Supplementary Information for

**Supplementary Figure 1.**
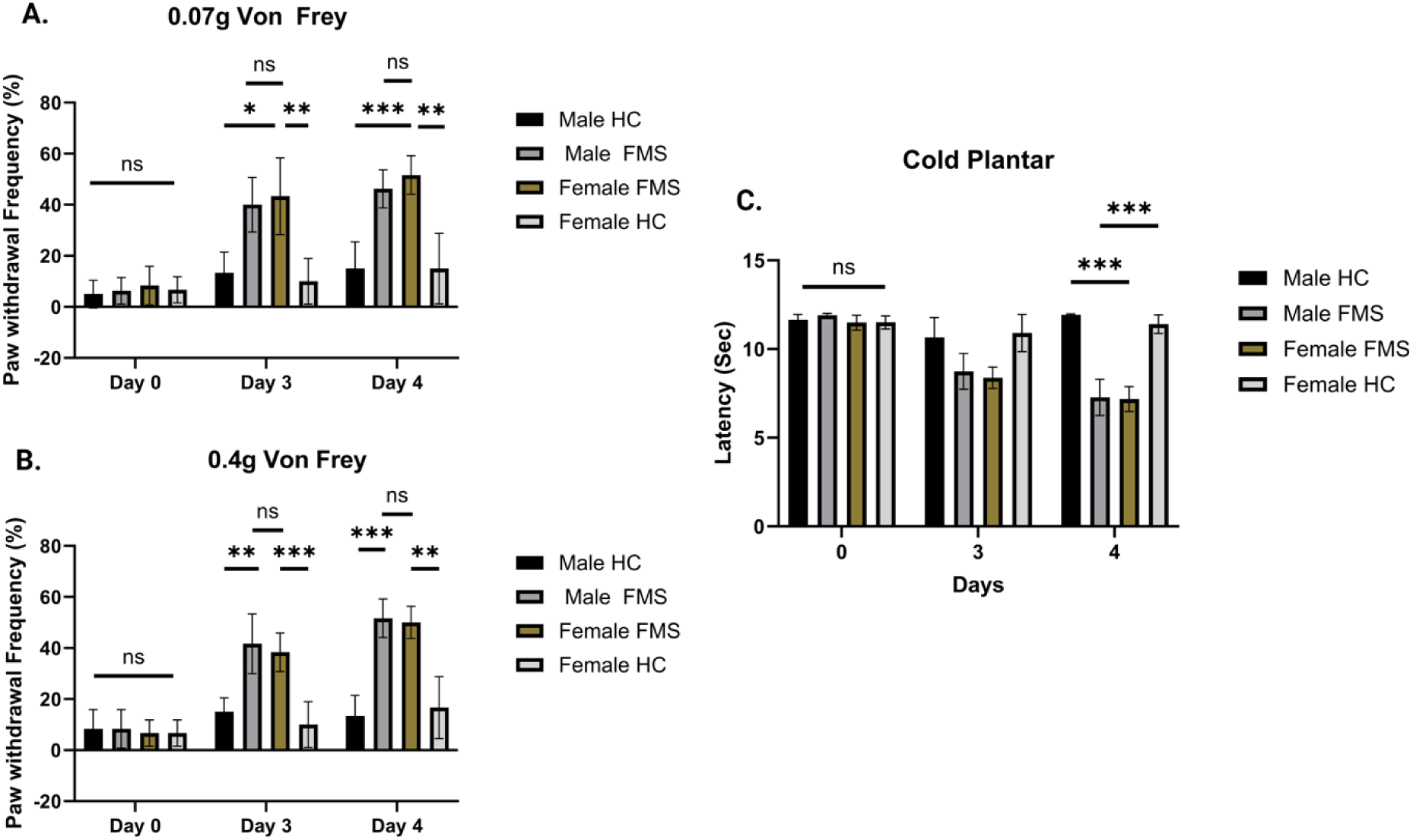
Male and Female Mice Show Comparable Responses to Fibromyalgia Syndrome Treatment with Human IgG Pain development is independent of the sex of the mice. Mice received intraperitoneal injections (IP) of 8mg IgG from fibromyalgia patients or healthy controls over 2 consecutive days from 3 different FMS patients or Healthy controls to evaluate whether IgG transfer triggers pain sensitivity. (A-B) Mechanical sensitivity was assessed using the von Frey test at low (0.07g) and high (0.4g) forces, with withdrawal responses from both hind paws shown. Data represent Male and Female groups (n = 6/group). (C) Cold allodynia was measured using the cold plantar assay, and responses from both hind paws Data represent Male and Female groups (n = 6/group). Statistical significance is indicated as ns = not significant, *p < 0.05, **p < 0.01, and ***p < 0.001. Data are presented as mean ± SEM and were analyzed using one-ANOVA followed by Bonferroni post hoc test. BioRender.com.

**Supplementary Figure 2.**
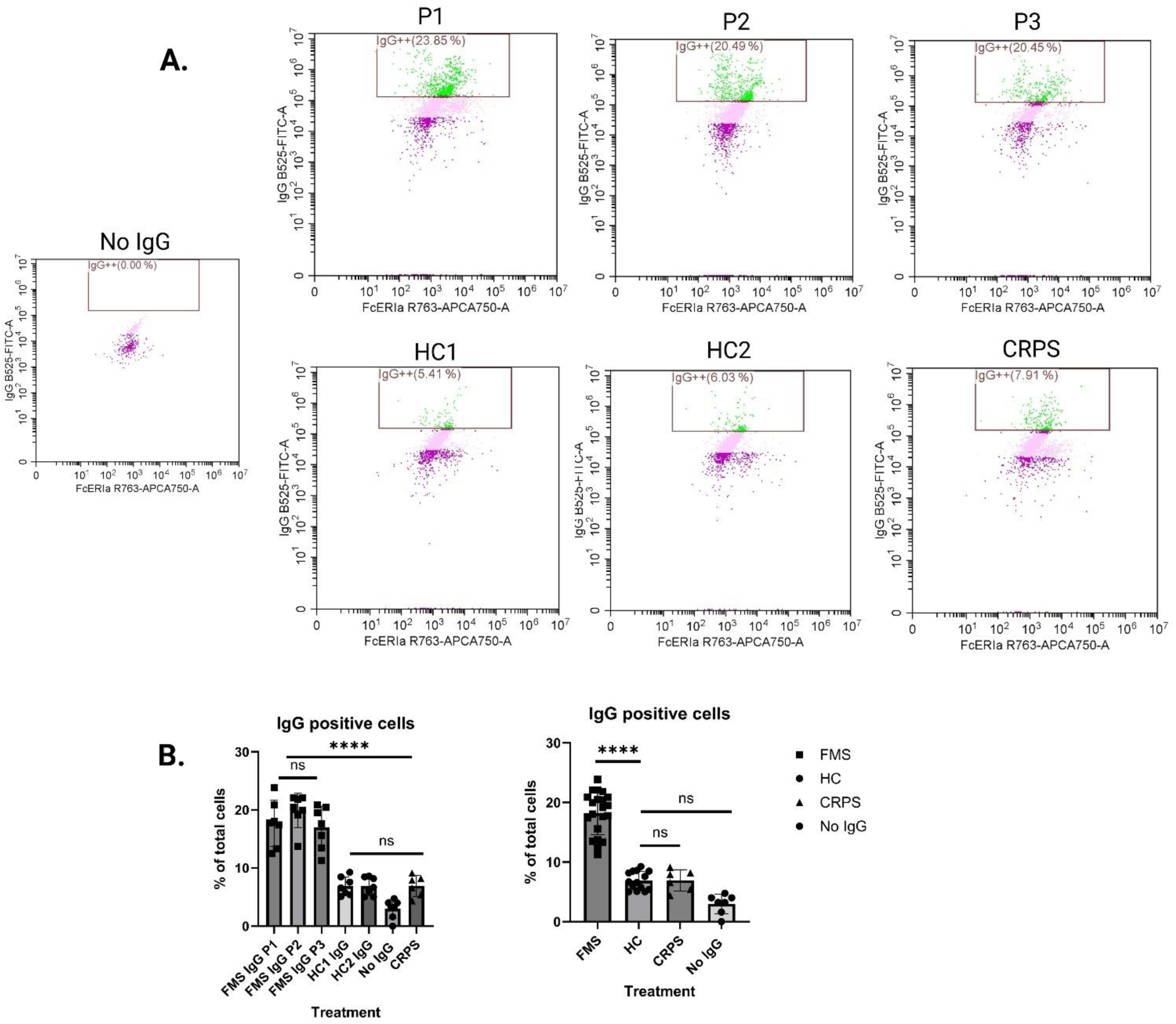
Binding Activity Comparison Between CRPS and FMS Samples Using flow cytometry, we assessed the binding of IgG from FMS patients and CRPS to mast cells (A) Representative flow cytometry panels showing the percentage of mast cells pre-gated for FcεRIα and C-kit, used to identify mast cells and the population positive for IgG. (B) Histogram representing the cell count of IgG binding in WT LAD2 mast cells. Comparisons between FMS IgG and CRPS IgG were analyzed using one-way ANOVA followed by Dunnett’s post-hoc test. ns = not significant; individual dots represent repeat experiments, *p < 0.05, **p<0.01 Created with BioRender.com.

**Supplementary Figure 3.**
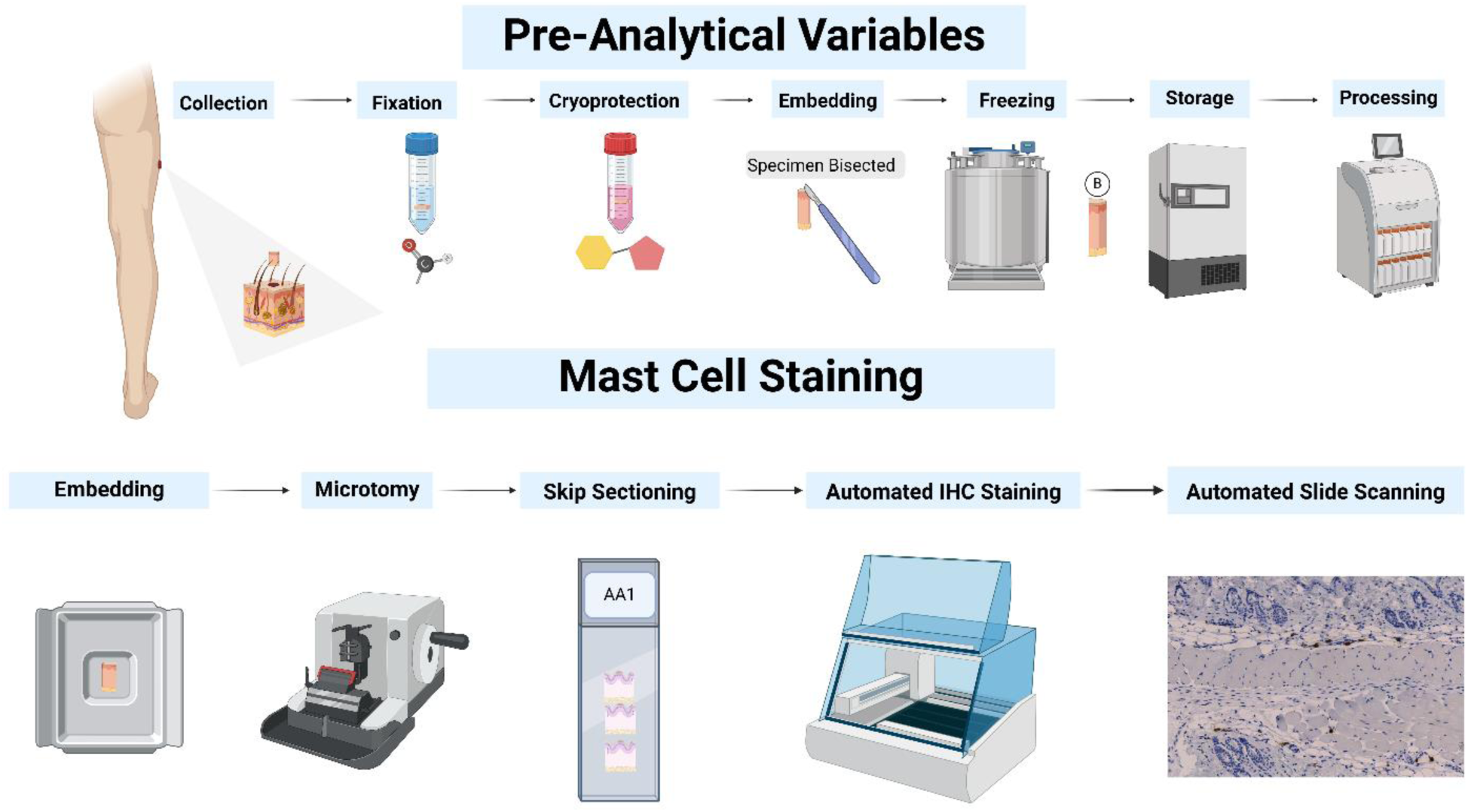
Workflow to Quantify Mast Cells in Human Skin using Automated Immunohistochemistry and Slide Scanning Skin tissue sections were processed from patients with FMS (n = 20) and controls (C) (n = 16) on an automated immunohistochemistry platform and visualized using a 3,3′-Diaminobenzidine (DAB) detection kit. The study quantified anti-tryptase 1 (AA1) positive mast cells within 4µm paraffin-embedded tissue sections. Sections were imaged using an automated slide scanner. QuPath (Version 0.5.1) was used to identify mast cells positively stained with hematoxylin and AA1 using DAB, following a standard research protocol for cell detection with a single gated threshold of 0.3 to categorize positive and negative cells. All statistical summaries and tests were conducted in R-studio (Version 4.0.2). The total detections per mm² (detections/area), total positive cells per mm² (DAB positive/area), and cumulative positive percentage (total positive detections x 100) were calculated.

**Supplementary Figure 4.**
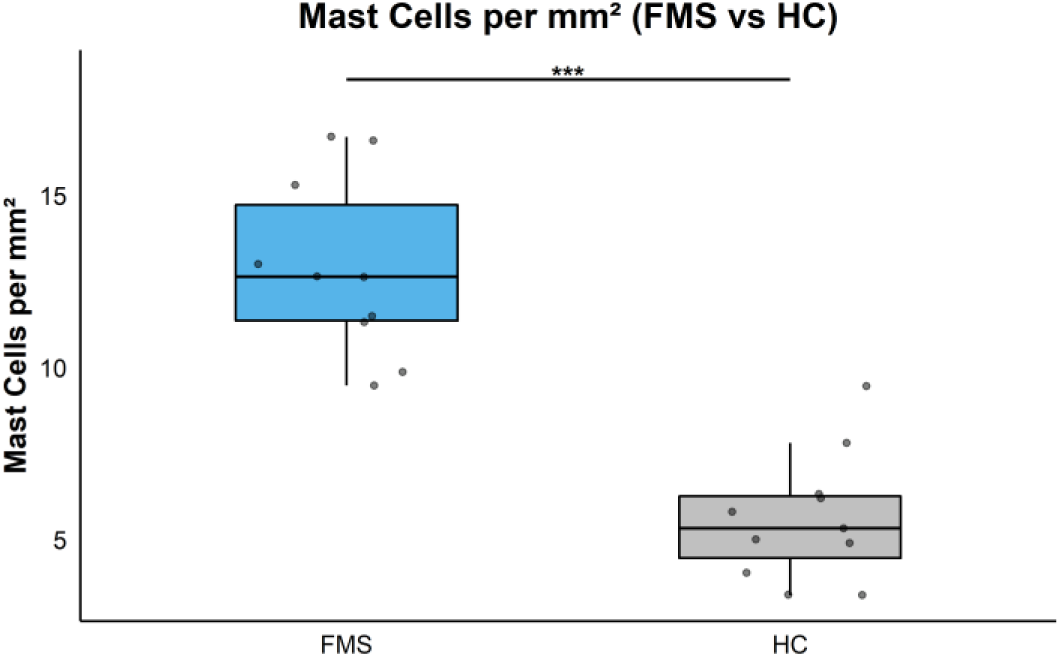
Increases mast cell density in non-glabrous mice skin following FMS treatment Using immunohistochemistry mast cells were detected using anti-tryptase antibodies visualized by DAB staining. Each data point represents reflects the analysis of one 4-micron tissue section. Mast cell density was significantly higher with FMS-IgG (FMS) relative to HC-IgG (HC) treated mice (12.9±2.6 vs 5.6±1.8 mast cells/mm²; mean difference +7.30 mast cells/mm²; ***p < 0.001).

**Supplementary Table 1:** Correlation Analysis of Phenotypic Symptoms and Mast Cell Density in Fibromyalgia Syndrome.

| Variable | Questionnaire Wording | R | R<sup>2</sup> | p-value | Test |
| --- | --- | --- | --- | --- | --- |
| Difficulty Urinating (SFNSL) | I have difficulties with urinating (either emptying my bladder or being able to hold my water) (Severity) | 0.52 | 0.27 | 0.02 | Spearman |
| Food Digestion Problems (SFNSL) | My food does not seem to go down well (Frequency) | 0.45 | 0.2 | 0.05 | Spearman |
| Pressure Sensitivity (PDQ) | Does slight pressure in this area, e.g., with a finger, trigger pain? | 0.41 | 0.17 | 0.08 | Spearman |
| Cold or Heat Sensitivity (PDQ) | Is cold or heat (bath water) in this area occasionally painful? | 0.39 | 0.16 | 0.09 | Pearson |
| Current Pain (PDQ) | How would you assess your pain now, at this moment? | 0.39 | 0.15 | 0.09 | Pearson |
| Bowel Problems (SFNSL) | I have problems with my bowel movements (Frequency) | 0.37 | 0.14 | 0.1 | Spearman |
| Dry Eyes (SFNSL) | I have dry eyes (Severity) | 0.37 | 0.14 | 0.11 | Spearman |
| PDQ Total | NA | 0.36 | 0.13 | 0.12 | Pearson |
| Average Pain (PDQ) | How strong was the pain during the past 4 weeks on average? | 0.33 | 0.11 | 0.15 | Pearson |
| SFNSL Total | NA | 0.33 | 0.11 | 0.16 | Pearson |
| Age | NA | 0.3 | 0.09 | 0.2 | Pearson |
| Burning Sensation (PDQ) | Do you suffer from a burning sensation (e.g., stinging nettles) in the marked areas? | 0.29 | 0.08 | 0.23 | Spearman |
| Palpitations (SFNSL) | I suffer from palpitations (Frequency) | 0.29 | 0.08 | 0.21 | Spearman |
| Painful Arms Seriousness (SFNSL) | I have painful arms (Severity) | 0.29 | 0.08 | 0.22 | Spearman |
| Strongest Pain (PDQ) | How strong was the strongest pain during the past 4 weeks? | 0.28 | 0.08 | 0.23 | Spearman |
| Numbness (PDQ) | Do you suffer from a sensation of numbness in the areas that you marked? | 0.26 | 0.07 | 0.28 | Pearson |
| Symptoms (FIQR) | NA | 0.26 | 0.07 | 0.27 | Pearson |
| Depression | NA | 0.22 | 0.05 | 0.36 | Spearman |
| Painful Arms Seriousness (SFNSL) | I have painful arms (Severity) | 0.22 | 0.05 | 0.35 | Spearman |
| McGill Pain Index | NA | 0.22 | 0.05 | 0.38 | Spearman |
| Blurred Vision (SFNSL) | I have blurred vision (Severity) | 0.21 | 0.04 | 0.39 | Spearman |
| Total (FIQR) | NA | 0.19 | 0.04 | 0.42 | Pearson |
| Chest Pain Seriousness (SFNSL) | I have chest pain (Severity) | 0.18 | 0.03 | 0.45 | Spearman |
| Cold Hands and Feet (SFNSL) | My feet and/or hands are colder than I am used to (Severity) | 0.18 | 0.03 | 0.44 | Spearman |
| Food Stuck (SFNSL) | I have the feeling that my food gets stuck in my throat (Severity) | 0.17 | 0.03 | 0.48 | Spearman |
| Function (FIQR) | NA | 0.15 | 0.02 | 0.53 | Pearson |
| Years Since Diagnosed with Fibromyalgia Syndrome | NA | 0.14 | 0.02 | 0.55 | Pearson |
| Chest Pain Frequency (SFNSL) | I have chest pain (Frequency) | 0.13 | 0.02 | 0.58 | Spearman |
| Tingling Sensation (PDQ) | Do you have a tingling or prickling sensation in the area of your pain (like crawling ants or electrical tingling)? | 0.1 | 0.01 | 0.68 | Spearman |
| Ocular Surface Disease Index Score | NA | 0.09 | 0.01 | 0.71 | Pearson |
| Cold Feet and Hands Frequency (SFNSL) | My feet and/or hands are colder than I am used to (Frequency) | 0.08 | 0.01 | 0.74 | Spearman |
| Pins and Needles in Hands | I have a tingling sensation in my hands (pins and needles) (Severity) | 0.08 | 0.01 | 0.75 | Spearman |
| Neuropathy Symptom Profile | NA | 0.07 | 0 | 0.77 | Pearson |
| Anxiety | NA | 0.06 | 0 | 0.81 | Spearman |
| Sheet Intolerance | At night I throw the bedclothes off my legs (Severity) | 0.06 | 0 | 0.81 | Spearman |
| McGill Pain Questionnaire Descriptors Total | NA | 0.04 | 0 | 0.85 | Pearson |
| Electric Shock Sensation | Do you have sudden pain attacks in the area of your pain, like electric shocks? | 0.04 | 0 | 0.86 | Spearman |
| Muscle Cramps | I suffer from muscle cramps (Frequency) | 0.03 | 0 | 0.91 | Spearman |
| Fibromyalgia Impact Questionnaire Revised Daily Life | NA | 0.01 | 0 | 0.97 | Pearson |
| Hot Flushes | I have sudden hot flushes (Severity) | 0.01 | 0 | 0.98 | Spearman |
| Light Touch Sensitivity | Is light touching (clothing, a blanket) in this area painful? | -<br>0.03 | 0 | 0.92 | Pearson |
| BMI | NA | -<br>0.03 | 0 | 0.9 | Spearman |
| Sensitive Skin Seriousness (SFNSL) | The skin of my legs is over-sensitive (Severity) | -<br>0.08 | 0.01 | 0.74 | Spearman |
| Pins and Needles in Legs (SFNSL) | I have a tingling sensation in my legs (pins and needles) (Severity) | -<br>0.14 | 0.02 | 0.56 | Spearman |
| Dizziness Seriousness (SFNSL) | I feel dizzy when I get up (Severity) | -<br>0.24 | 0.06 | 0.3 | Spearman |
| BMI – Body Mass Index; FIQR – Revised Fibromyalgia Impact Questionnaire; PDQ – Pain Detect Questionnaire; SFNSL – Small Fibre Neuropathy Screening List; |  |  |  |  |  |

**Supplementary Table 2:** commercial antibodies, provide supplier name, catalogue number.

| Anigen/Reagent | Fluorochrome | Species reactivity | Vendor | Catalog number | Dilution |
| --- | --- | --- | --- | --- | --- |
| CD117 (c-kit) | BV421 | Human | BioLegend | 313215 | 1:200 |
| FcεRIα | APC-Cy7 | Human | BioLegend | 334631 | 1:200 |
| CD45 | PE | Human | STEMCELL | 60018PE.1 | 1:200 |
| Avidin | Texas Red | Human/mouse | Millipore | A2348-5MG | 1:500 |
| Anti-IgG | FITC | Human | Thermo Fisher | A-11013 | 1:200 |
| Claudin-1 | FITC | Human/mouse | Santa Cruz | sc-166338 | 1:500 |

